# The role of prooxidants and antioxidants in shaping life-history and parasite tolerance in *Anopheles* mosquitoes

**DOI:** 10.1101/2024.12.17.628837

**Authors:** Tiago G. Zeferino, Alfonso Rojas Mora, Jacob C. Koella

**Affiliations:** Institute of Biology, University of Neuchâtel, Rue Emile-Argand 11, 2000 Neuchâtel, Switzerland

**Author notes:** Corresponding author (TGZ).

## Abstract

Oxidative homeostasis plays important roles in physiology, for reactive oxygen species not only lead to damaging oxidative stress, but also regulate important physiological processes like immunity and longevity. ROS are therefore expected to be a key factor underlying many host-parasite interactions. We evaluated the role of the host’s oxidative status on the outcome of infection with the mosquito *Anopheles gambiae* infected by the microsporidian *Vavraia culicis*. To do so, we manipulated the oxidative status of the mosquitoes by feeding them early (the first four days after emergence) or late (from five days after emergence onwards) either a standard sugar source or one supplemented with a prooxidant (hydrogen peroxide) or antioxidant (vitamin C), and then measured the longevity and fecundity of uninfected and infected mosquitoes and (for infected mosquitoes) the parasite load at a given day (13 days after emergence) or when the mosquitoes died. The prooxidant generally increased longevity, but if consumed early after emergence its impact was lessened by the infection by *Vavraia*. In contrast the antioxidant increased fecundity, and the impact was not affected by the status of infection or by the timing of consumption. Finally, early consumption of both supplements increased *Vavraia*’s spore load at 13 days after emergence and at death. In contrast, late consumption enhanced the parasite’s growth late in the mosquito’s life. Thus, our experiment revealed complex effects of prooxidant and antioxidant consumption, emphasising the critical role of timing and context in shaping their influence on biological traits.

## Introduction

Nutrition is a crucial factor shaping host-parasite interactions because of the energy it delivers to the host and the parasite and because of micronutrients and other compounds that affect the parasite’s growth ^1–4^ and fitness ^5^. One mechanism is that nutrition modulates constitutive and inducible immune responses ^2,6–12^. However, since nutrition influences immunity through various direct and indirect pathways, including the interactions with the host’s endogenous microbiota ^3,13,14^, many details of the mechanism are poorly understood ^15,16^. Yet, a deeper understanding of the relationship between nutrition and immunity is important, as it has significant implications for morbidity and mortality ^2,17–20^.

For many animals a large part of the nutrition is provided by flowering plants. Since these have coevolved with their pollinators and herbivores for millions of years ^21^, they contain numerous volatile and non-volatile compounds ^22^ in their nectar, pollen, resin and oils ^23,24^. These compounds include macronutrients (proteins, fats and carbohydrates). While the macronutrients ^7,8,10,12,25,26^ influences the outcome of host-parasite interactions directly and by interacting with each other ^6,9,11^, their impact on immune function and the host’s survival is complicated by other plant-derived compounds such as micronutrients and secondary metabolites, which may also interfere with the host-parasite interaction.

Micronutrients are chemical substances like vitamins and minerals that are required in small quantities for normal growth, development, and the functioning of the immune system ^22^. Several phytochemicals in nectar or pollen act as antimicrobials agents that protect not only the plants and their flowers from pathogens ^27–29^, but also the animals that feed on them ^30–32^. Such medicinal properties may be particularly significant for pollinators and nectar-feeders, which are frequently exposed to the secondary metabolites.

As regular nectar-feeders ^33^ and as vectors for parasites with major implications for human health, mosquitoes are particularly interesting for studying effects of secondary metabolites in nectar on host-parasite interactions. Indeed, nectar influences the outcome of mosquito infections by malaria parasites, likely due to a combination of toxic secondary metabolites and the nutritional profile of the nectar ^34^. Many of the compounds in nectar possess oxidative or antioxidative properties, which impact the production of reactive oxygen species (ROS) and the development of oxidative stress (OS) in mosquitoes ^35^. Both ROS and OS help to shape life-history traits of mosquitoes ^36–38^ and to regulate their immune responses ^39^. Indeed, redox cycles induced by prooxidants like hydrogen peroxide and antioxidants like ascorbic acid have been suggested as the most likely defence mechanism against a broad spectrum of bacteria and fungi in nectar ^40,41^. However, while these protective mechanisms help mosquitoes defend against parasitic infections, other, damaging consequences of oxidative stress imposes selective pressure on mosquitoes to finely balance their oxidative homeostasis. Understanding this balance would offer insights into how mosquito nutrition affects their role in the transmission of human diseases. Furthermore, understanding the roles of macro- and micronutrients in oxidative homeostasis and host-parasite interactions would help to clarify the behavioural ecology of self-medication ^31,42,43^.

In a recent study, we found that *Anopheles gambiae* mosquitoes altered their dietary preferences in response to an infection by the microsporidian parasite *Vavraia culicis* ^44^. Infected mosquitoes selectively consumed a prooxidant or an antioxidant, with their choices varying based on the infection stage. By contrast, uninfected mosquitoes actively avoided these diets. This selective intake influenced the oxidative homeostasis of mosquitoes, supporting the idea of self-medication and suggesting potential effects on other life-history traits.

Here, we sought to further explore the broader impacts of dietary supplementation with a prooxidant or an antioxidant on mosquitoes’ health and their possible implications for disease transmission. We thus examined the effects of consuming a standard sugar source or one supplemented with a prooxidant or an antioxidant, administered early or late in life, on *A. gambiae* longevity, fecundity, and resistance to *V. culicis* infection. Additionally, we assessed whether these supplements impose a cost on uninfected mosquitoes.

## Materials and Methods

### Experimental system

We used the Kisumu strain of *A. gambiae s*.*s* ^45^, which had been maintained at our standard laboratory conditions (about 600 individuals per cage, constant access to 6% sucrose solution, 26 ± 1ºC, 70 ± 5% relative humidity and 12 h light/dark) for many years before the experiments.

To infect mosquitoes, we used the microsporidian *V. culicis floridensis*, which had been provided by J.J Becnel (USDA, Gainesville, FL, USA). *V. culicis* is an obligate intracellular parasite of several mosquito genera, including *Aedes, Culex, Anopheles, Culiseta, Ochlerotatus* and *Orthopodomyia* ^46–48^. To maintain its status as a generalist parasite, we alternately infected *Aedes aegypti* and *A. gambiae* ^49^.

Mosquitoes become infected as larvae when they ingest spores. After several rounds of replication, the parasite produces infectious spores that spread to the gut and fat body cells. These spores are released into larval habitats when larvae die, when adults die on the water surface, or when eggs coated with spores are laid onto the water.

### Mosquito rearing and maintenance

We conducted two experiments, one for longevity and one for fecundity, in a way that ensures that the two traits do not interfere with each other due to the life-history trade-off in *A. gambiae* ^50^. The design of the two experiments was identical.

Freshly hatched (0 to 3-hour-old) larvae were individually placed into 12-well culture plates, each containing 3 ml of deionised water. The larvae received Tetramin Baby® fish food every day according to their age (0.04, 0.06, 0.08, 0.16, 0.32 and 0.6 mg/larva, respectively, ate ages 0, 1, 2, 3, 4 and 5 or older ^51^).

Two-day-old larvae were exposed to either 0 or 10,000 spores of *V. culicis*. Pupae were transferred to individual 50 ml falcon tubes with approximately 10 ml of deionised water ^50^. Upon emergence, females were maintained according to the experiment: for the longevity-experiment we kept females alone in 150 ml cups to prevent mating, whereas in the fecundity-experiment we kept mosquitoes in groups of 20 females and 20 males in 1.5 L jars to allow mating (see details below).

### Dietary treatments

We gave the mosquitoes one diet (referred to as early diet) from day 0 (the day of emergence) to day 4, and another diet (referred to as late diet) from day 5 up to the mosquitoes’ death. This categorization was based on our previous work ^44^ where we showed that infected mosquitoes exhibited a preference for different diets in early and late stages of life.

At each of these stages, mosquitoes received either 10% sucrose (referred to as sugar diet), 10 % sucrose supplemented with 8mM of hydrogen peroxide (prooxidant diet) or 10% sucrose supplemented with 1mg/mL of ascorbic acid (antioxidant diet). The diets were freshly prepared on the day they were given to the mosquitoes. The concentrations of hydrogen peroxide and ascorbic acid correspond to concentrations found in nectar ^52–55^, and they had been tested before the experiment to ensure they did not kill the mosquitoes.

### Longevity experiment

Adult females were individually moved to 150 ml cups (5.5 cmØ x 10 cm) covered with a net. The cups contained 50 ml deionised water covered by a petri dish (50 mmØ) to prevent the mosquitoes from drowning. A 10 × 7 cm filter paper partly submerged in the water maintained humidity in the cup. The mosquitoes were provided with cotton balls soaked in the diet they were allocated to. The cotton balls were replaced daily. The cups were surveyed daily, and dead individuals were collected and frozen at - 20ºC for later analysis.

### Fecundity experiment

Females were moved 1.5 L jars (15 cm Ø x 20 cm) in groups of 20 according to their treatment status. To enable mating, we also added 20 males. These were from our colony and were thus not infected so that fecundity was not influenced by the infection of their mates. The mosquitoes were fed daily with cotton balls soaked in their respective diet. Nine days after emergence females were given the opportunity to blood-feed on TGZ’s arm for five minutes. One day later females that were not fully engorged were excluded from the experiment. The next day the remaining females were transferred to individual cups containing 50 ml deionised water covered by a petri dish (50 mm Ø) to prevent the mosquitoes from drowning. The cup also contained a 94 mm Ø filter paper folded into the shape of a cone for egg laying. Four days after blood-feeding (so 13 days after emergence), the females were frozen at -20ºC for later analysis and a picture was taken of each filter paper so that we could count the eggs.

### Measurements

The number of eggs was measured from the photos with the software ImageJ v1.54 ^56^.

The right wing of every mosquito was removed. Its length was measured from the axillary incision to the wing’s tip ^57,58^ with ImageJ.

The remainder of the mosquito was put into a 2 ml Eppendorf tube containing 0.1 ml of deionised water. A stainless-steel bead (Ø 5 mm) was added to each tube and the samples were homogenised with a Qiagen TissueLyser LT at a frequency of 30 Hz for two minutes. The number of spores in each sample was counted from 0.1 µl of the sample with a haemocytometer under a phase-contrast microscope (400x magnification).

### Statistical analysis

All analyses were conducted with R ^59^ version 4.4.1, using the packages DHARMa ^60^, car ^61^, glmmTMB ^62^, emmeans ^63^ and multcomp ^64^. Significance was assessed with the Anova function of the car package ^61^. We used a type III ANOVA in the case of a significant interaction and a type II ANOVA otherwise. When relevant, we performed post-hoc multiple comparisons with the package emmeans, using the default Tukey adjustment.

Age at death was analysed with a linear model with a Gaussian distribution of errors, where the response variable was age at death and the explanatory variables were infection status, early diet, late diet, and their interactions. Wing length was included as a covariate.

The proportion of individuals laying eggs was analysed with a generalised linear model with a binomial distribution of errors, where the response variable was the proportion of individuals that laid eggs and the explanatory variables were the infection status, early diet, late diet, and their interactions. Wing length was included as a covariate.

The number of eggs was analysed with a linear model with a Gaussian distribution of errors, where the response variable was egg count and the explanatory variables were infection status, early diet, late diet, and their interactions. Wing length was included as a covariate. Individuals that did not lay eggs were excluded from this analysis.

Mosquitoes with detectable spores were used to assess the spore load. Because we counted the number of spores in a haemocytometer containing 0.1 µl of the sample (i.e. 1/1000 of the total volume), the detection threshold was estimated to be 1000 spores. Spore load on day 13 (from the fecundity experiment) was analysed using a generalised linear model with a negative binomial distribution of errors ^65^, where the response variable was spore load and the explanatory variables were early diet, late diet, their interaction, and wing length as a covariate. Since the number of spores increases throughout the mosquitoes’ life, we analysed the spore load at death (from the longevity experiment) using a generalised linear model with a negative binomial distribution of errors, where the response variable was spore load at death and the explanatory variables were age at death, early diet, late diet, their interaction, and wing length as a covariate.

## Results

### Longevity

Infection by *V. culicis* shortened the lifespan from an average of 29.4 days to 24.8 days (χ^2^ = 83.11, df = 1, p < 0.001) and feeding on sugar supplemented with the prooxidant increased the lifespan, whether the consumption was early (prooxidant: 29.9 days; antioxidant: 26.2 days; sugar: 25.1 days; χ^2^ = 65.75, df = 2, p < 0.001) or late (prooxidant: 28.1 days; antioxidant: 26.9 days; sugar: 26.2 days; χ^2^ = 8.94, df = 2, p = 0.011, **Fig. 2**). However, the impact of late diet was small, and its impact was not seen within groups defined by early diet and infection status (multiple comparisons of **Fig. 2**). The effects of the early and the late diet were independent of each other (early diet * late diet: χ^2^ = 2.69, df = 4, p = 0.611). However, the early diet had a stronger impact on uninfected individuals than infected ones, with both supplemented diets increasing longevity (prooxidants: 32.7 vs 27.1 days; antioxidants: 29.1 vs 23.3 days; sugar 26.3 vs. 23.9 days; interaction infection status * early diet: χ^2^ = 10.11, df = 2, p = 0.006, **Fig. 2ab**), while the impacts of the late diet (infection status * late diet: χ^2^ = 0.96, df = 2, p = 0.617) and of the combination of the two diets (infection status * early diet * late diet: χ^2^ = 1.62, df = 4, p = 0.805) on infected and uninfected individuals were similar.

**Figure 1.**
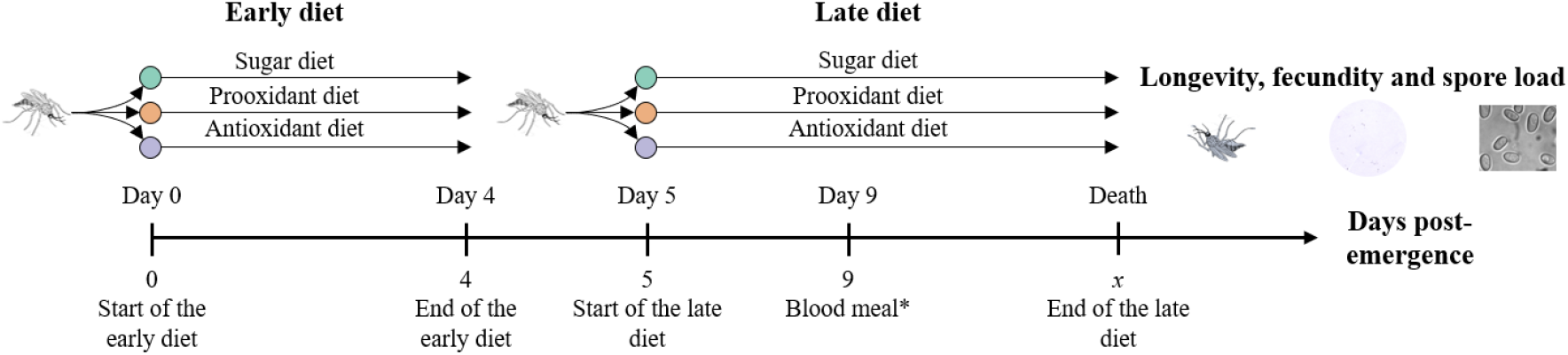
Experimental design. Illustration of the dietary treatments and experimental setup in this study. Longevity and spore load at death were measured during the longevity experiment, while fecundity and spore load on day 13 were assessed during the fecundity experiment. * Indicates the blood meal taken exclusively by individuals in the fecundity experiment.

**Figure 2.**
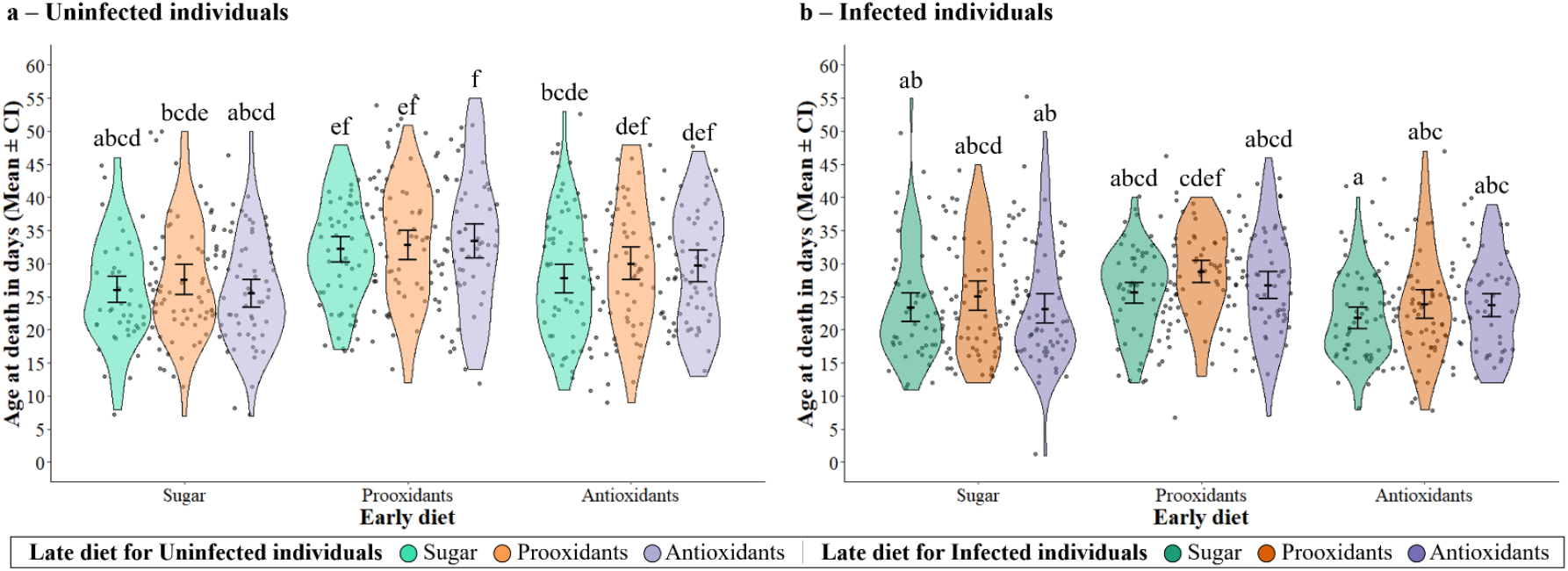
Age at death of **(a)** uninfected and **(b)** infected mosquitoes according to the diet given early and late in life. The samples sizes were 60 individuals per diet and infection status. The error bars show the 95% confidence intervals of the means and the letters indicate statistically significant differences from multiple comparisons.

### Fecundity

59.5% of the mosquitoes laid at least one egg, independently of the their infection status (57.8% vs 61.0%, χ^2^ = 1.12, df = 1, p = 0.290), early diet (62.3% vs 56.9% vs 59.0%, χ^2^ = 2.13, df = 2, p = 0.345), late diet (61.2% vs 60.1% vs 57.0%, χ^2^ = 1.25, df = 2, p = 0.536) or any of their interactions (infection status * early diet: χ^2^ = 3.95, df = 2, p = 0.139; infection status * late diet: χ^2^ = 1.76, df = 2, p = 0.414; early diet * late diet: χ^2^ = 5.91, df = 4, p = 0.206; infection status * early diet * late diet: χ^2^ = 2.64, df = 4, p = 0.619; **Fig. 3ab**).

**Figure 3.**
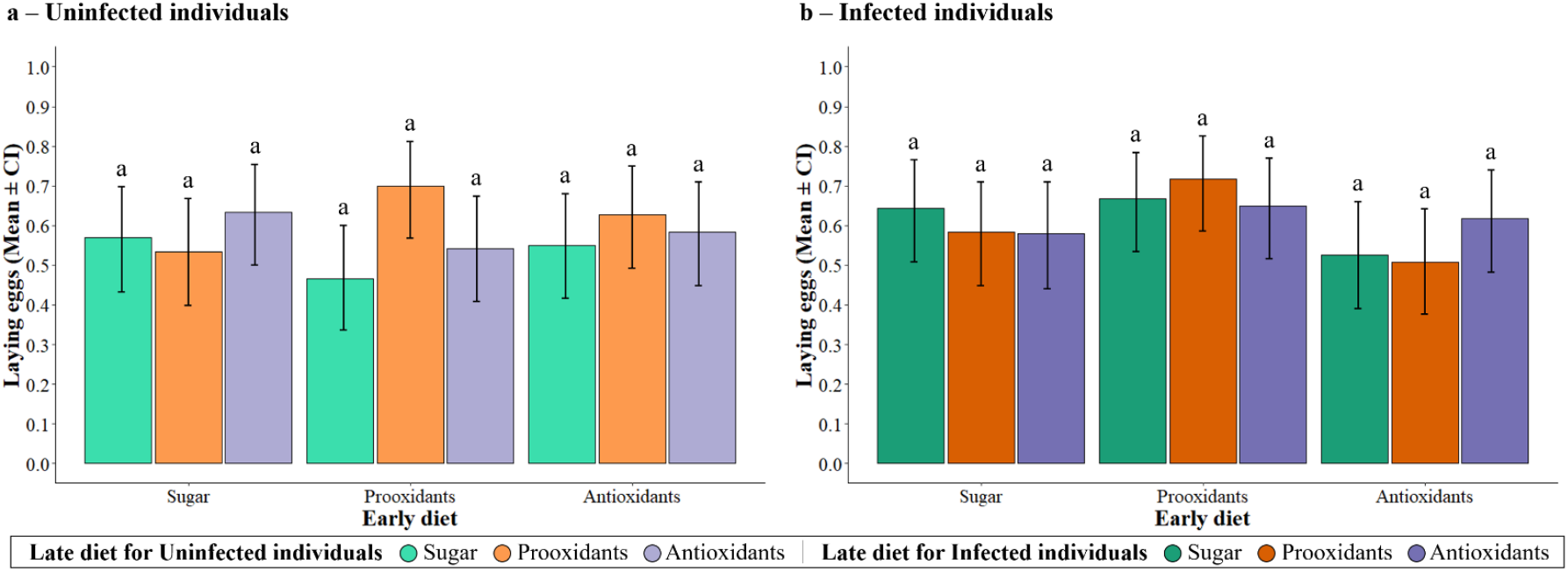
The proportion of **(a)** uninfected and **(b)** infected individuals that laid eggs according to the diet given early and late in life. The sample sizes were between 58 and 60 individuals in **(a)** and between 57 and 60 individuals in **(b)**. The error bars show the 95% confidence intervals of the mean proportions among jars and the letters indicate statistically significant differences from multiple comparisons.

In contrast, if the mosquitoes did lay eggs, infection by *V. culicis* reduced the number of eggs from an average of 87.5 eggs to 81.7 eggs (χ^2^ = 4.06, df = 1, p = 0.044) and consuming sugar supplemented with either a prooxidant or, in particular, an antioxidant increased the number of eggs, whether the supplement was given early (prooxidant: 85.9 eggs; antioxidant: 95.3 eggs; sugar: 73.6 eggs; χ^2^ = 23.91, df = 2, p < 0.001) or late (prooxidant: 89.4 eggs; antioxidant: 87.8 eggs; sugar: 76.9 eggs; χ^2^ = 13.06, df = 2, p = 0.001). These effects were independent of any combination of the treatments (infection status * early diet: χ^2^ = 3.66, df = 2, p = 0.160; infection status * late diet: χ^2^ = 2.55, df = 2, p = 0.280; early diet * late diet: χ^2^ = 4.42, df = 4, p = 0.352; infection status * early diet * late diet: χ^2^ = 5.62, df = 4, p = 0.229; **Fig. 4ab**).

**Figure 4.**
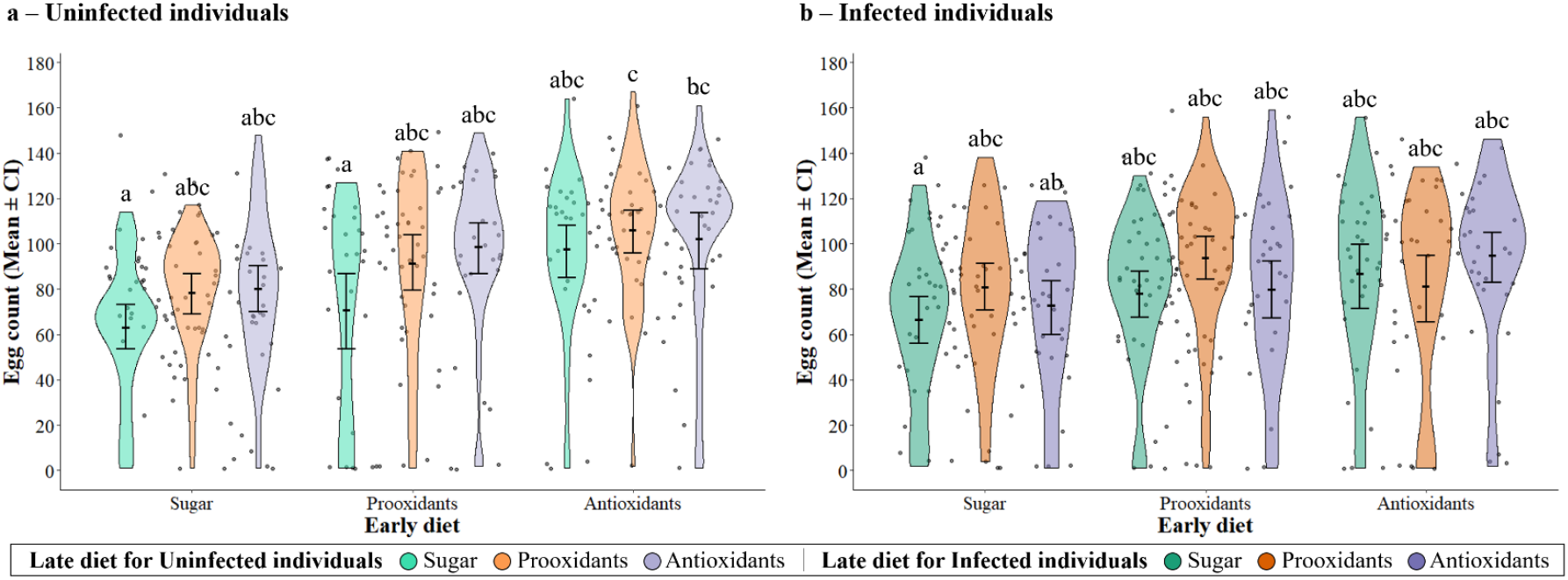
Number of eggs laid by **(a)** uninfected and **(b)** infected individuals according to the diet given early and late in life. The sample sizes were 33, 31, 38, 28, 42, 32, 33, 37 and 35 in **(a)**, and 38, 35, 33, 40, 43, 39, 30, 30 and 37 in **(b)**. The error bars show the 95% confidence intervals of the means and the letters indicate statistically significant differences from multiple comparisons.

### Infection dynamics

The mosquitoes that were assayed 13 days after emergence harboured, on average, about 1.5×10^5^ spores (**Fig. 5a)**, and early consumption of sugar supplemented with either a prooxidant or an antioxidant increased the spore load (prooxidant: 1.7×10^5^ spores; antioxidant: 1.6×10^5^ spores; sugar: 1.1×10^5^ spores; χ^2^ = 8.35, df = 2, p = 0.015). However, although spore load was not influenced by their late diet (prooxidant: 1.4×10^5^ spores; antioxidant: 1.6×10^5^ spores; sugar: 1.3×10^5^ spores; χ^2^ = 2.31, df = 2, p = 0.315), early consumption of prooxidants followed by late consumption of antioxidants led to the higher parasite load (early diet * late diet: χ^2^ = 12.15, df = 4, p = 0.016).

**Figure 5.**
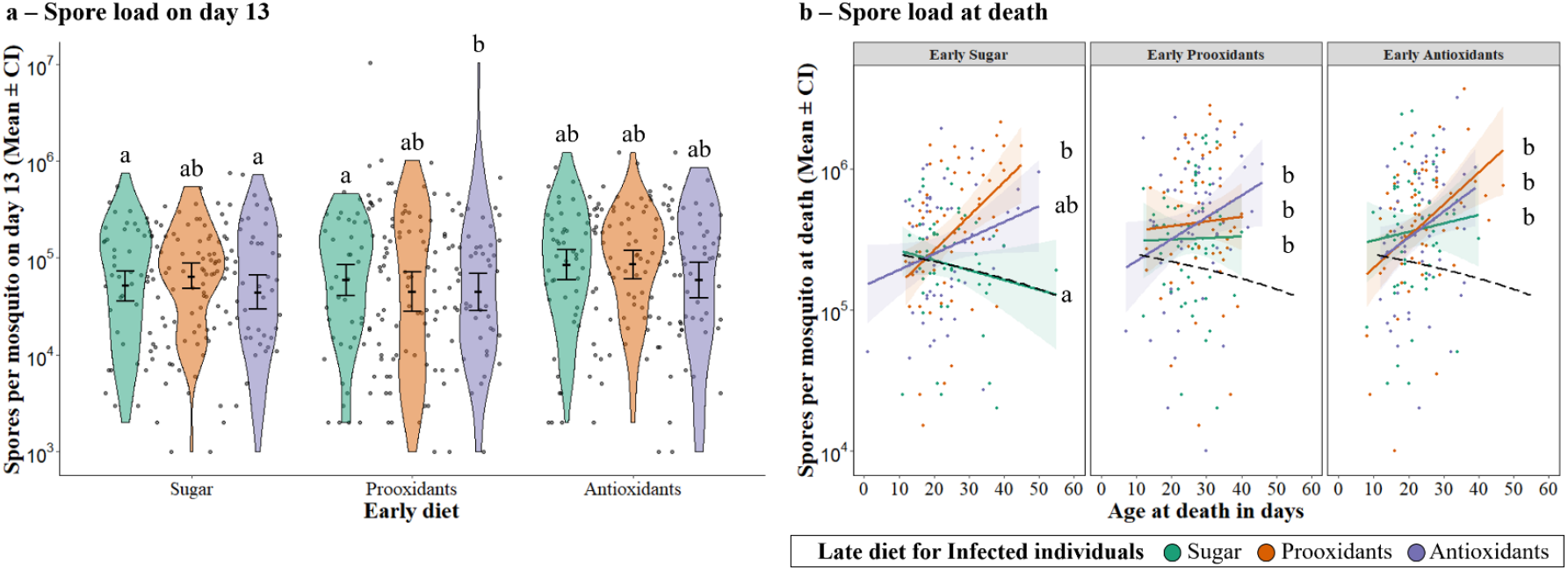
Spore load on day 13 **(a)** and spore at death **(b)** according to the diet given early and late in life. The samples sizes were 60 individuals per diet and infection status. The error bars show the 95% confidence intervals of the means and the letters indicate statistically significant differences from multiple comparisons. A smooth line was drawn for each treatment group using the “lm” method in ggplot2 to illustrate the relationship between spore load at death (log10 scale) and age at death. The shaded regions represent 95% confidence intervals around the regression lines. A dashed black line represents this relationship for individuals taking the sugar diet early and late.

The spore load at death increased with age at death (χ^2^ = 25.40, df = 1, p < 0.001) and was higher in mosquitoes consuming either supplemented diet provided early (χ^2^ = 25.30, df = 2, p < 0.001, **Fig. 5b**), with an average spore load of 5.9×10^5^ per day in mosquitoes feeding on the prooxidant and antioxidant diet, and 4.1×10^5^ spores on the sugar diet. Notably, while the late diet had a modest effect on spore load (χ^2^ = 9.10, df = 2, p = 0.010), it also changed the relationship between spore load and age, with supplemented diets increasing spore load with age at death (age * late diet: χ^2^ = 9.36, df = 2, p = 0.009, **Fig. 5b**).

## Discussion

Supplementing the sugar meals of mosquitoes with prooxidants or antioxidants had significant effects on their longevity and fecundity and on parasite growth, likely increasing the mosquitoes’ reproductive success. These effects, however, varied depending on the timing of the supplementation, with early diets having a particularly strong influence. In the following sections, we delve deeper into our results, offering a possible explanation for how these dietary treatments may benefit infected and uninfected mosquitoes, discuss their connections to oxidative homeostasis and dietary preferences ^44^, and consider the broader epidemiological implications of these findings.

### Early Dietary Supplementation Enhances Mosquito Longevity and Reproductive Output

Consuming a prooxidant or an antioxidant together with sugar, particularly early in life, significantly extended the mosquitoes’ longevity (**Fig. 2**) and fecundity (**Fig. 4**). The beneficial effect of antioxidants aligns with findings in other studies. For instance, antioxidants enhance longevity in mealworm beetles by suppressing immune activity and reducing immunopathology ^66^. Similarly, feeding crickets with vitamin C reduces several aspects of oxidative damage, including damaged DNA in haemocytes ^67^. The positive effect of prooxidants observed in our study, however, contrasts with an experiment in which *Anopheles arabiensis* fed with hydrogen peroxide live less long. This discrepancy may be due to the low, naturally occurring dose of hydrogen peroxide used in our study. At such levels, ROS from the diet may exhibit beneficial effects, such as enhancing viral defence mechanisms ^68^ or modulating insulin-signalling pathway to extend longevity ^69^.

### Parasitic Infection Modulates the Benefits of Dietary Supplements Through Oxidative Stress

The effects of dietary supplementation were also observed in infected mosquitoes, though to a lesser extent. Two main explanations may account for this difference. First, parasites may reduce the beneficial impacts of the supplements. For antioxidants, this could occur because parasitic infections stimulate the immune system, leading to oxidative damage. Although antioxidant intake typically reduces oxidative stress and limits cellular damage ^31,70^, infection-induced immune activation might counteract these benefits. For prooxidants, the combined ROS increase from the diet and parasite-induced immune stimulation may push OS beyond manageable levels, disrupting oxidative homeostasis and leading to oxidative damage.

Second, these dietary supplements may reduce the pathological effects of infection on longevity and fecundity, although while increasing parasite load (**Fig. 5**). This increase, observed both at day 13 and at death, suggests that the diets help hosts to better cope with the physiological costs of infection, enabling them to live despite a more efficiently parasite proliferation. This aligns with a form of tolerance where the host is not reducing the parasite’s growth (as in resistance) but is instead managing the infection’s impact on its own survival (longevity-tolerance ^71–74^), fecundity (fecundity-tolerance ^71,75– 78^) and potentially other life-history traits.

Although both diets disrupt the mosquito’s oxidative homeostasis ^44^, their effects diverged. Prooxidants may boost immune function, perhaps by raising ROS levels, which helps to combat infections ^79,80^, and increases longevity (**Fig. 2**). In contrast, antioxidants appeared to suppress immune responses by decreasing ROS ^81–85^. While this boosted fecundity (**Fig. 4**), it shortened lifespan slightly and compromised the ability to contain infections, highlighting the immuno-suppressant effects of antioxidants diets.

### Timing of consumption suggests different mechanisms by which these diets have a benefit

While these dietary items did not clear the infection, as is often the case in the wild ^86,87^, they significantly improved the overall fitness of the host, as reflected in increased longevity and fecundity ^32,42,88,89^. Notably, the timing of supplementation played a critical role, with early diet interventions showing stronger effects than late supplementation. This temporal difference aligns with the infection dynamics of *V. culicis*, which undergoes exponential growth up to day four before slowing and reaching a plateau around day twelve ^49^.

Our findings suggest that our dietary treatments are most effective when administered before day five, during the early, rapid replication phase of the parasite. Prooxidant intake during this period likely increases oxidative stress, which, while causing some cellular damage ^78,90,91^, may also inhibit *V. culicis* replication, thus extending host longevity ^70,75^. In contrast, antioxidant intake reduces oxidative stress, mitigating cellular damage but potentially facilitating *V. culicis* replication. This could explain the observed trade-off between increased fecundity and shorter lifespan in mosquitoes consuming antioxidants ^31,70^. These results emphasise the importance of the interaction between dietary timing and the physiological dynamics of both the host and the parasite.

### The complex interplay between prooxidants and antioxidants challenges the traditional notion of oxidative stress

Our findings underscore the complex interplay between prooxidants and antioxidants in mosquito physiology, challenging the traditional notion of oxidative stress as solely damaging. Instead, oxidative stress appears to play a dual role, where its benefits and costs are carefully balanced by the mosquitoes, influencing their survival and reproduction. This nuanced perspective highlights the adaptive strategies mosquitoes employ to regulate oxidative stress in response to dietary factors, emphasizing the importance of oxidative balance in their biology.

Understanding this complexity provides valuable insights into mosquito ecology and physiology, offering potential avenues for innovative vector control strategies. However, our results also raise important considerations for the use of *V. culicis* as a biological control agent. Infected mosquitoes showed an ability to mitigate the fitness costs of parasitism, potentially reducing the effectiveness of microsporidians. These findings emphasise the need to account for host-parasite dynamics and dietary influences when designing biological control programs.

## Supporting information

Supplementary information for "The role of prooxidants and antioxidants in shaping life-history and parasite tolerance in Anopheles mosquitoes"

## Acknowledgements

We thank Luís M. Silva for his advice and technical support. TGZ and the project were supported by SNF grant 310030_192786.

## Author contributions

ARM conceived the overall idea. TGZ, ARM and JCK designed the experiments. TGZ collected, analysed, and interpreted the data and wrote the first draft of the manuscript. All authors contributed critically to the drafts.

## Data availability

All data generated or analysed during this study are included as Supplementary Information files.

## Additional Information

The authors declare no competing interests.

