## Supplementary information for "The role of prooxidants and antioxidants in shaping life-history and parasite tolerance in Anopheles mosquitoes" for "The role of prooxidants and antioxidants in shaping life-history and parasite tolerance in *Anopheles* mosquitoes"

**Table S1.** Age at death.

**Table S2.** Proportion of individuals laying eggs.

**Table S3.** Egg count.

**Table S4.** Spore load.

**Table S1. Age at death.**

| Tested effect | | | |
| --- | --- | --- | --- |
| **Model 1** – Age at death | **df** | $\chi^2$ | ***p*** |
| Infection status | 1 | 83.11 | **<0.001** |
| Early | 2 | 65.75 | **<0.001** |
| Late | 2 | 8.94 | **0.011** |
| Wing length | 1 | 14.56 | **<0.001** |
| Infection status:Early | 2 | 10.11 | **0.006** |
| Infection status:Late | 2 | 0.96 | 0.617 |
| Early:Late | 4 | 2.69 | 0.611 |
| Infection status:Early:Late | 4 | 1.62 | 0.805 |

**Table S2. Proportion of individuals laying eggs.**

| **Tested effect** | | | |
|---|---|---|---|
| **Model 2** – Proportion of individuals laying eggs | **df** | $\chi^2$ | ***p*** |
| Infection status | 1 | 1.12 | 0.290 |
| Early | 2 | 2.13 | 0.345 |
| Late | 2 | 1.25 | 0.536 |
| Wing length | 1 | 0.28 | 0.597 |
| Infection status:Early | 2 | 3.95 | 0.139 |
| Infection status:Late | 2 | 1.76 | 0.414 |
| Early:Late | 4 | 5.91 | 0.206 |
| Infection status:Early:Late | 4 | 2.64 | 0.619 |

**Table S3. Egg count.**

| Tested effect | | | |
|---|---|---|---|
| **Model 3** – Egg count | **df** | **$\chi^2$** | ***p*** |
| Infection status | 1 | 4.06 | **0.044** |
| Early | 2 | 23.91 | **<0.001** |
| Late | 2 | 13.06 | **0.001** |
| Wing length | 1 | 12.49 | **<0.001** |
| Infection status:Early | 2 | 3.66 | 0.160 |
| Infection status:Late | 2 | 2.55 | 0.280 |
| Early:Late | 4 | 4.42 | 0.352 |
| Infection status:Early:Late | 4 | 5.62 | 0.229 |

**Table S4. Spore load.**

| Tested effect | | | |
|---|---|---|---|
| **Model 4a** – Spore load on day 13 | **df** | $\chi^2$ | ***p*** |
| Early | 2 | 8.35 | **0.015** |
| Late | 2 | 2.31 | 0.315 |
| Wing length | 1 | 7.31 | **0.007** |
| Early:Late | 4 | 12.15 | **0.016** |
| **Model 4b** – Spore load at death | **df** | $\chi^2$ | ***p*** |
| Age | 1 | 25.40 | **<0.001** |
| Early | 2 | 25.30 | **<0.001** |
| Late | 2 | 9.10 | **0.010** |
| Wing length | 1 | 10.65 | **0.001** |
| Age:Early | 2 | 1.23 | 0.541 |
| Age:Late | 2 | 9.36 | **0.009** |
| Early:Late | 4 | 6.73 | 0.151 |
| Age:Early:Late | 4 | 7.43 | 0.114 |
